# Deciphering state bridges via a single-cell large language model

**DOI:** 10.64898/2026.11.21.768275

**Authors:** Qintong Fei, Maodi Liang, Chunmei Cui, Boming Kang, Qinghua Cui

**Author notes:** To whom the correspondence should be addressed: Boming Kang, Qinghua Cui.

## Abstract

Large-scale single-cell transcriptomic data provide a new avenue for precision medicine by uncovering cellular heterogeneity. However, systematic comparison of remodeling patterns across various biological states remains unavailable. Here, we first developed a computational framework based on a foundation model of single cell biology, Universal Cell Embedding (UCE), to harmonize annotations comprising 36 cell types. Based on an integrated dataset of single cell RNA sequencing (scRNA-seq) with 8,789 human samples comprising more than 59 million cells or nuclei across 614 biological states, we then embedded each state with a 36-dimensional vector in a unified space of cell-composition change and further deciphered states bridges by analyzing the relationships among the 614 state vectors. As a result, 22,161 significant bridges (P < 0.05) between the 614 states were revealed. Moreover, we successfully identified a number of well-known bridges such as bridge between Alzheimer’s disease and Parkinson’s disease (P = 2.21e-15), suggesting the effectiveness of the proposed framework. More importantly, we uncovered a number of new bridges. Furthermore, analyses of the number of changed cell types (NCC) and Shannon entropy revealed heterogeneous and tissue-dependent patterns of cellular composition remodeling. This study provides a new strategy for identifying disease associations and potential therapeutic relationships by deciphering bridges between biological states from scRNA-seq landscape.

## 1 Introduction

In recent years, single-cell transcriptomic studies have characterized changes in cellular composition across a wide range of diseases and treatment states^[1, 2]^. These studies have shown that inflammatory, fibrotic, and malignant states are often accompanied by extensive cellular remodeling, including immune-cell infiltration, stromal-cell activation, and parenchymal-cell depletion^[3–5]^. These studies paved a new avenue for precision medicine by exploring cellular dynamics. However, these dynamics have largely been interpreted independently within individual datasets or disease contexts and therefore remain disconnected from one another. Consequently, although we have an increasingly detailed understanding of which cell populations change in a given state, it remains difficult to determine whether two seemingly unrelated states undergo similar or even opposing patterns of cellular remodeling. This gap limits our ability to uncover potential biological relationships beyond existing knowledge.

Encoding complex biological states into comparable features and using these features to uncover relationships between states has precedents at the molecular level. The Connectivity Map (CMap) introduced the use of gene-expression signatures to connect diseases, drugs, and other perturbations^[6]^. This paradigm was subsequently expanded by large-scale expression-profile compendia^[7]^ and resources such as CREEDS^[8]^, LINCS^[9]^, and iLINCS^[10]^, which progressively incorporated feature information from multiple omics layers. These paradigms and resources have greatly advanced the development of precision medicine. Nevertheless, existing signature-matching approaches capture molecular changes, such as alterations in gene or protein expression level, but cannot directly represent how different cell populations are remodeled. This raises an important question: can changes in cellular composition serve as a multicellular remodeling signature and be used to decipher the potential bridges among various biological states?

The above large-scale comparisons of various biological states require a unified representation of cell identity. However, single-cell studies differ substantially in sequencing platforms, data-processing pipelines, and cell-annotation systems. The same cell population is often assigned different names or annotated at different levels of granularity, limiting direct comparisons of cellular composition across datasets^[11, 12]^. The recently developed Universal Cell Embedding (UCE) single-cell foundation model maps cells from different tissues and datasets into a shared embedding space, which provides a foundation model for harmonizing cell identities across heterogeneous single-cell datasets and establishing a unified cell-type framework^[13]^.

Here, we developed a single-cell analytical framework for deciphering bridges across various biological states. We first integrated large-scale scRNA-seq data and used UCE to establish a unified annotation system comprising 36 cell types. We then represented each state as a 36-dimensional vector, thereby mapping the states into a shared space of cell-composition change. On this basis, we deciphered state bridges by analyzing relationships among the 614 state vectors. We further quantified the number of changed cell types and Shannon entropy to characterize the extent and evenness of cellular composition remodeling and to investigate their variation across tissue sources. The framework not only uncovered state bridges supported by established biological knowledge but, more importantly, also revealed previously unrecognized state bridges. This study proposed a new paradigm for discovering disease associations and potential therapeutic relationships from large-scale scRNA-seq landscapes.

## 2 Materials and Methods

### 2.1 Data collection and quality control

Human single-cell transcriptomic data were primarily obtained from the DISCO and scCT-DB databases^[14, 15]^, with additional datasets from HeartMap and the human brain aneurysm single-cell atlas^[16, 17]^. Publicly available scRNA-seq and single-nucleus RNA sequencing (snRNA-seq) datasets released up to June 2026 were collected. Datasets were included if they met the following criteria: (1) samples were of human origin; (2) tissue origin, disease or intervention group, and matched control information were clearly defined; and (3) cell-composition information and essential metadata were available. Datasets involving enrichment or sorting for specific cell types, lacking a clearly defined control group, or missing essential metadata were excluded. Routine purification procedures for cells or nuclei used only to remove debris, doublets or multiplets were not considered exclusion criteria. After screening and deduplication, a total of 8,789 samples comprising 59,056,428 cells or nuclei were retained for downstream analyses.

Cell-level quality control was performed using Scanpy version 1.10.2^[18]^. Cells were retained if they contained more than 100 detected genes, more than 300 total UMI counts, and less than 30% mitochondrial transcripts. Doublets were identified using Scrublet version 0.2.3^[19]^, with the expected doublet rate set to 5%. At the gene level, low-abundance genes were removed, and only genes detected in at least three cells were retained.

### 2.2 Unified cell-type annotation based on UCE

Universal Cell Embedding (UCE) is a Transformer-based foundation model for single-cell transcriptomics that was self-supervised on large-scale, multi-species and multi-tissue single-cell transcriptomic data. By integrating protein embeddings with deep encoding, UCE maps individual cells into a unified 1,280-dimensional representation and enables zero-shot embedding across datasets without additional fine-tuning^[13]^. In this study, UCE embeddings were computed for each sample independently, with parallel computation performed across four NVIDIA A800 GPUs.

Tabula Sapiens v2 was used as the reference dataset for cell-type annotation. This dataset contains expert-curated human single-cell data from multiple tissues, with standardized cell-type annotations and high biological reliability^[13, 20]^. UCE embeddings were first generated for cells in the reference dataset, and a multiclass logistic regression classifier was trained using the corresponding standardized cell-type labels. UCE embeddings were then computed for all cells in the study cohort, and cell types were predicted using the trained classifier. Finally, all cells were harmonized into 36 cell types for downstream cell-composition analyses.

### 2.3 Construction of cell-composition change profiles

Changes in cellular composition were quantified according to a dataset–sample–state hierarchy. Each independent dataset represented a single-cell study with a relatively distinct experimental design and sequencing workflow. A sample represented a profiled specimen or sequencing library; some donors contributed multiple samples. A biological state was defined as a group of samples within the same dataset that shared the same disease, physiological condition or intervention. All cell-composition changes were calculated within individual datasets, using matched controls from the same dataset as the baseline to minimize the influence of technical variation across datasets.

For each sample, we first calculated the proportion of each of the 36 cell types relative to the total number of cells. Cell-type proportions were then averaged across all samples belonging to the same state to generate a cell-composition profile for that state. Using the corresponding matched control group within the same dataset as the baseline, fold changes (FC) in cell-type proportions were calculated. A pseudocount of 1.00e-6 was added to avoid division-by-zero errors caused by zero values. Finally, each biological state was represented as a 36-dimensional feature vector comprising the proportional changes of the 36 cell types.

### 2.4 Cross-state bridge analysis

To compare patterns of cellular-composition remodeling across various biological states, we used the 36-dimensional cell-composition change vector of each state as its feature representation and performed cross-state association analyses using Spearman rank correlation, cosine similarity and Fisher’s exact test. Firstly, Spearman rank correlation coefficients (Rho) and corresponding P values were calculated between pairs of states to assess concordance in cell-type change patterns. Meanwhile, cosine similarity was calculated between the 36-dimensional log2-transformed FC (log_2_FC) vectors to quantify similarity in the overall direction of cellular-composition changes. In addition, cell types were classified as increased (FC > 1.3) or decreased (FC < 1/1.3) according to predefined FC thresholds. For each pair of states, contingency tables were constructed according to the directions of cell-type changes, and Fisher’s exact test was used to calculate the odds ratio (OR) and corresponding P value, thereby assessing the association between the directions of cell-proportion changes in the two states.

### 2.5 Statistical analysis and visualization

Statistical analyses and data visualization were primarily performed using Python version 3.10.20 and R version 4.2.0. Both state-level UMAP and heatmaps of cell-composition changes were generated using the log_2_FC profiles of the 36 cell types. Hierarchical clustering for the heatmaps was performed using Euclidean distance. For state-level UMAP analysis, the number of nearest neighbors (n_neighbors) was set to 15 and the minimum distance (min_dist) to 0.1. A representative subset of cells was additionally constructed for cell-level UMAP visualization. First, 30% of all samples were randomly selected without replacement. For each cell type, up to 100,000 cells were retained; when fewer than 100,000 cells were available, all cells of that type were retained. The resulting representative subset comprised 2,611,156 cells or nuclei. UMAP dimensionality reduction was then performed using the 1,280-dimensional UCE embeddings of these representative cells, with n_neighbors set to 15 and min_dist set to 0.5.

Spearman correlations between all pairs of biological states were calculated using the change profiles of the 36 cell types. Hierarchical clustering was performed using 1 − Rho as the distance metric, and the resulting pairwise correlations were visualized as a correlation heatmap. State pairs with P < 0.05 and |Rho| > 0.65 were retained to construct the correlation network. After removal of isolated nodes, the core association network was extracted using a k-core threshold of 2.

## 3 Results

### 3.1 A large-scale scRNA-seq resource enables cross-state comparisons

The overall study design and analytical workflow are shown in **Figure 1**. Systematic comparison of cellular remodeling across biological states requires a sufficiently broad data foundation. To this end, we integrated human scRNA-seq data from DISCO, scCT-DB, and other independent public resources (**Figure 2A**). The resulting cohort comprised 8,789 samples and more than 59 million cells or nuclei across 614 biological states, providing a large-scale single-cell resource for cross-state comparisons.

**Figure 1.**
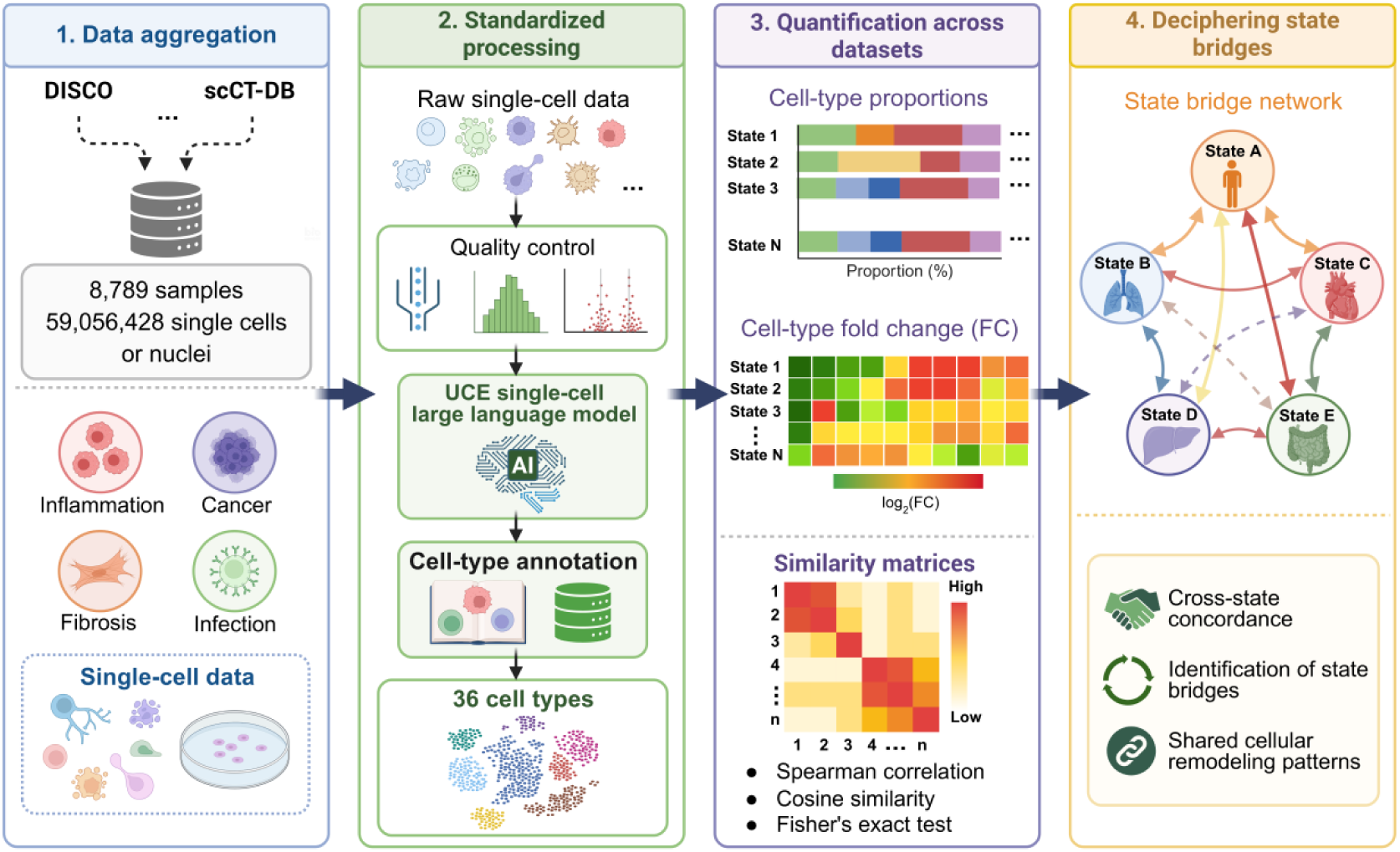
Study design and workflow. Public single-cell datasets were aggregated and subjected to standardized quality control and UCE-based cell-type annotation. Cell-type proportions and fold changes were then quantified across biological states, followed by cross-state similarity analyses using Spearman correlation, cosine similarity and Fisher’s exact test. These analyses were used to identify state bridges and shared cellular remodeling patterns across diseases and interventions.

**Figure 2.**
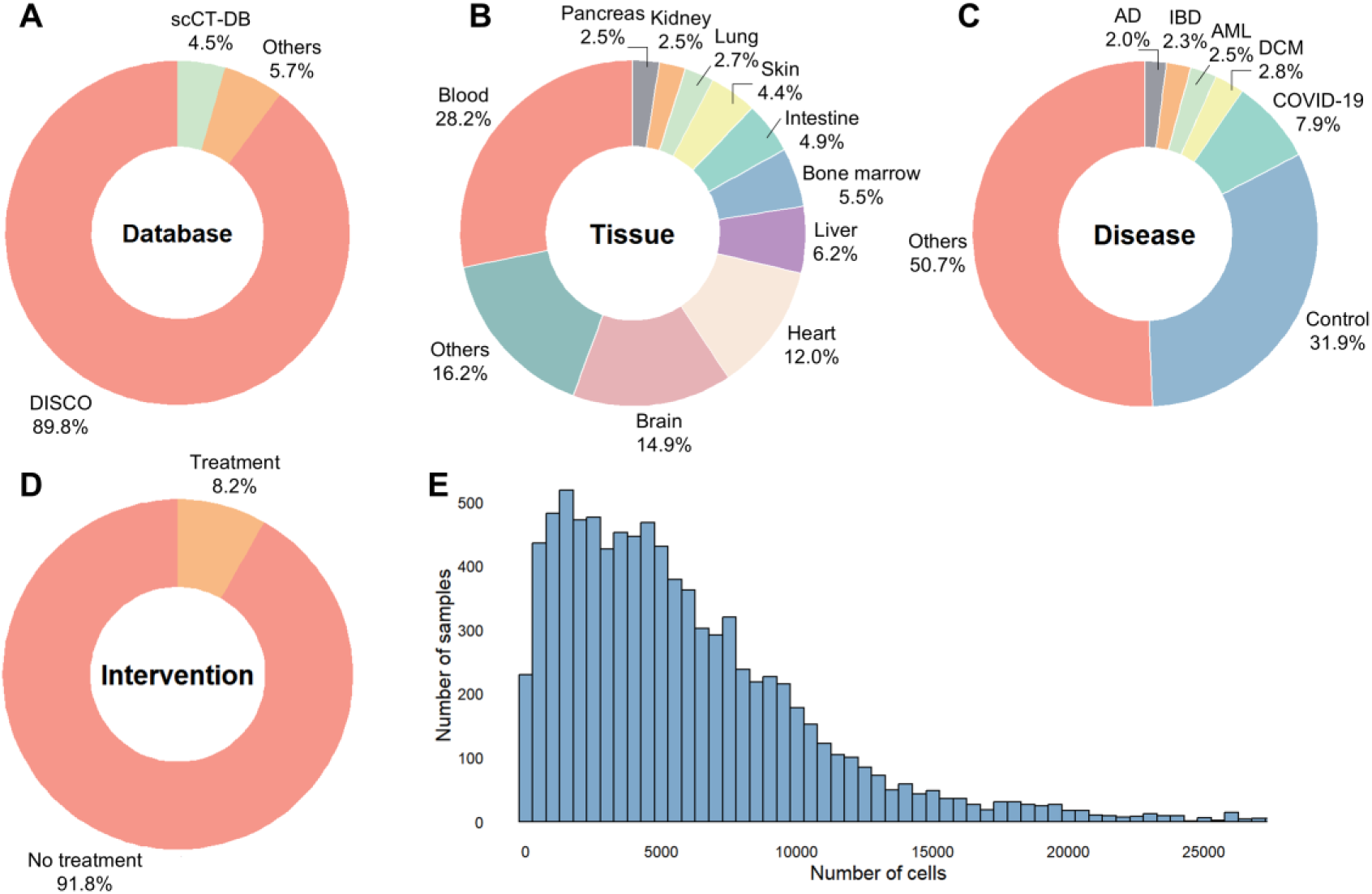
Overview of the sample composition of the large-scale single-cell study cohort. **(A)** Distribution of data sources. **(B)** Distribution of tissue types. **(C)** Distribution of disease and control groups. **(D)** Distribution of intervention status. **(E)** Histogram showing the distribution of cell numbers per sample.

The cohort covered a broad range of tissue origins and pathological contexts. Samples were derived from multiple tissues, including blood, brain, heart, and liver. Blood and brain accounted for the largest proportions, representing 28.2% and 14.9% of all samples, respectively (**Figure 2B**). The disease states encompassed a wide range of diseases and other pathological conditions, including coronavirus disease 2019 (COVID-19), acute myeloid leukemia (AML), and Alzheimer’s disease (AD) (**Figure 2C**). In addition, 8.2% of the samples were derived from treatment or intervention conditions (**Figure 2D**). After quality control, most samples contained between 3,000 and 8,000 cells (**Figure 2E**), indicating a relatively consistent distribution of cell numbers across samples. Overall, this broad coverage of tissues, diseases, and interventions enables different biological states to be analyzed within a common framework, extending comparisons beyond individual diseases or datasets.

### 3.2 UCE establishes a shared cell representation across states

A key prerequisite for large-scale cross-state comparisons is that the same cell types from different studies can be mapped to a consistent cell-identity framework. To achieve this, we used the Universal Cell Embedding (UCE) model^[13]^ to generate unified representations for the ∼59 million human cells and mapped them to 36 cell types. These cell types spanned major immune, neural, stromal, and epithelial lineages. T cells, myeloid leukocytes, neurons, and glial cells were among the most abundant cell types in the integrated cohort (**Figure 3A**). We next performed UMAP dimensionality reduction on UCE embeddings from a representative subset of cells. When colored by cell type, cells formed relatively distinct clusters in the low-dimensional space (**Figure 3B**). In contrast, cells from different datasets were broadly intermingled (**Figure 3C**). These results indicate that the unified UCE representation primarily captures cell identity rather than dataset origin.

**Figure 3.**
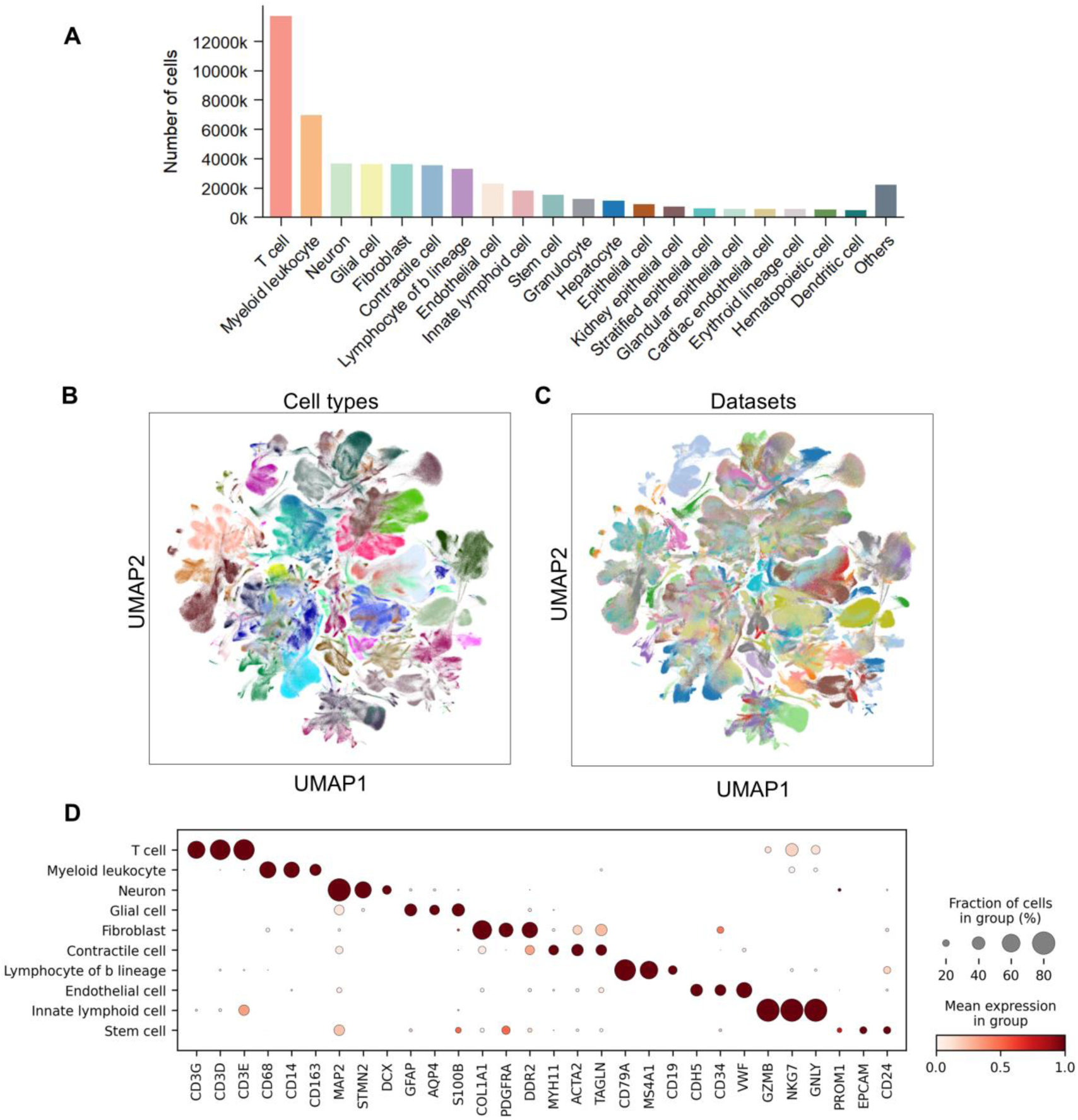
Unified cell-type annotation based on UCE. **(A)** Bar plot showing the distribution of cell numbers across the 20 most abundant cell types; lower-abundance cell types were grouped as “Others”. **(B)** UMAP visualization of cells based on UCE embeddings, colored by cell type. **(C)** The same UMAP embedding colored by dataset of origin, showing the distribution of cells from different datasets in the UCE embedding space. **(D)** Dot plot showing the expression of representative marker genes across the 10 most abundant cell types. Dot size indicates the proportion of cells expressing each gene within the corresponding cell type, and color indicates the average expression level of that gene within the corresponding cell type.

We further validated the annotations using canonical marker genes. T cells showed high expression of the T-cell markers CD3D and CD3E, whereas myeloid leukocytes expressed myeloid markers such as CD14 and CD68. Neurons showed high expression of neuronal markers, including MAP2 and STMN2 (**Figure 3D**). Overall, each cell type exhibited marker-gene expression patterns consistent with its known biological identity, supporting the accuracy of the unified cell-type annotations. Together, UCE established a common cell-identity reference across heterogeneous scRNA-seq datasets, enabling cells from different studies to be compared within a consistent cell-type framework.

### 3.3 Cellular remodeling features define a unified state space

After establishing a unified cell-type reference, we next asked whether different biological states could be transformed into directly comparable features. For each of the 614 states, we used the matched control from the same dataset as the baseline and calculated changes in the proportions of the 36 cell types. These changes were integrated into a 36-dimensional vector for each state (Supplementary Table 1) and mapped into a shared cell-composition change space. The heatmap of cell-composition changes showed that the 614 states exhibited highly heterogeneous remodeling patterns (**Figure 4A**), with distinct changes across multiple cell types. Importantly, these patterns were not driven by changes in a single cell population. Instead, each state was characterized by coordinated changes across multiple cell types, providing a unified representation for comparing remodeling patterns across diverse biological states.

**Figure 4.**
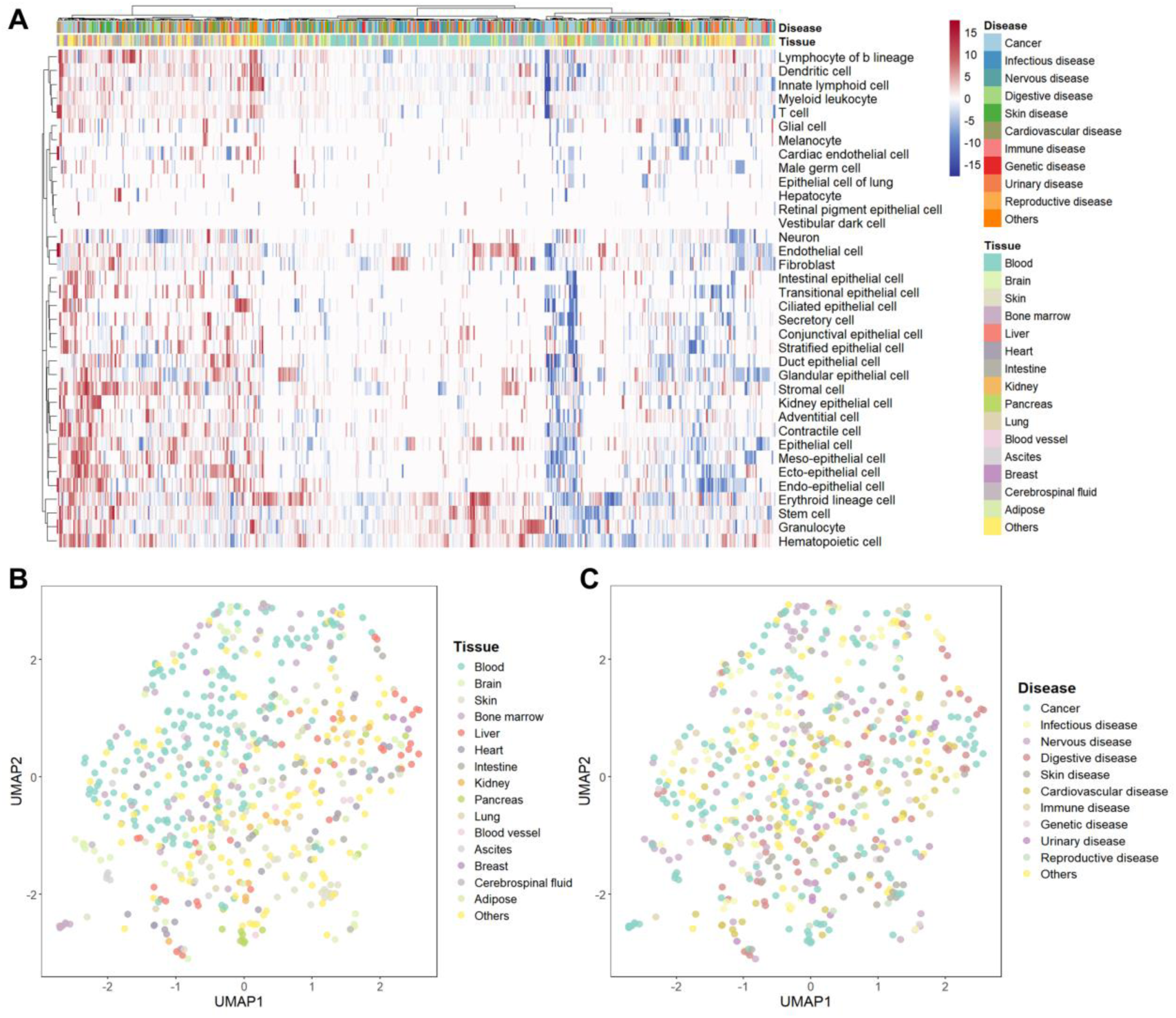
Cell-composition change profiles define a unified representation space across biological states. **(A)** Heatmap of cell-composition changes across 614 biological states. Columns represent biological states and rows represent cell types. Colors indicate changes in cell-type proportions relative to matched controls (log_2_FC), with red and blue denoting increases and decreases, respectively. Top annotations indicate the disease category and tissue of origin for each state. **(B)** UMAP of biological states constructed from cell-composition change profiles and colored by tissue of origin. **(C)** The same UMAP colored by disease category.

We next reduced the dimensionality of the 614 state vectors to examine how this representation relates to conventional classification frameworks. States derived from different tissues were broadly intermingled in the low-dimensional state space and did not form clear tissue-specific clusters (**Figure 4B**). Similarly, states from different disease categories did not segregate into distinct clusters according to conventional disease classifications (**Figure 4C**). These observations indicate that tissue origin or disease category does not fully determine the pattern of cellular-composition remodeling. States from different tissues or disease categories can still exhibit similar cellular changes. Thus, the 36-dimensional cellular-remodeling vectors not only distinguish the remodeling characteristics of individual states but also provide a common coordinate system that extends across tissue and disease labels. Conventional disease classifications describe what these states are, whereas this state space enables us to ask whether different states undergo cellular remodeling in similar ways. This shift from categorical labels to remodeling features provides a basis for identifying biological relationships beyond existing classifications.

### 3.4 The cellular-remodeling space reveals widespread state bridges

In the unified cellular-composition remodeling space, states from different tissues and disease contexts were broadly intermingled, suggesting potential relationships beyond existing biological knowledge. To systematically characterize these relationships, we analyzed the 36-dimensional cell-composition change vectors of the 614 biological states. Spearman correlation analysis identified 22,161 significant state bridges (P < 0.05), including 11,947 positive bridges and 10,214 inverse bridges (**Figure 5A**). The correlation heatmap and network analysis further revealed extensive connections among states, characterized by similar or opposing patterns of cellular-composition remodeling (**Figure 5B, 5C**).

**Figure 5.**
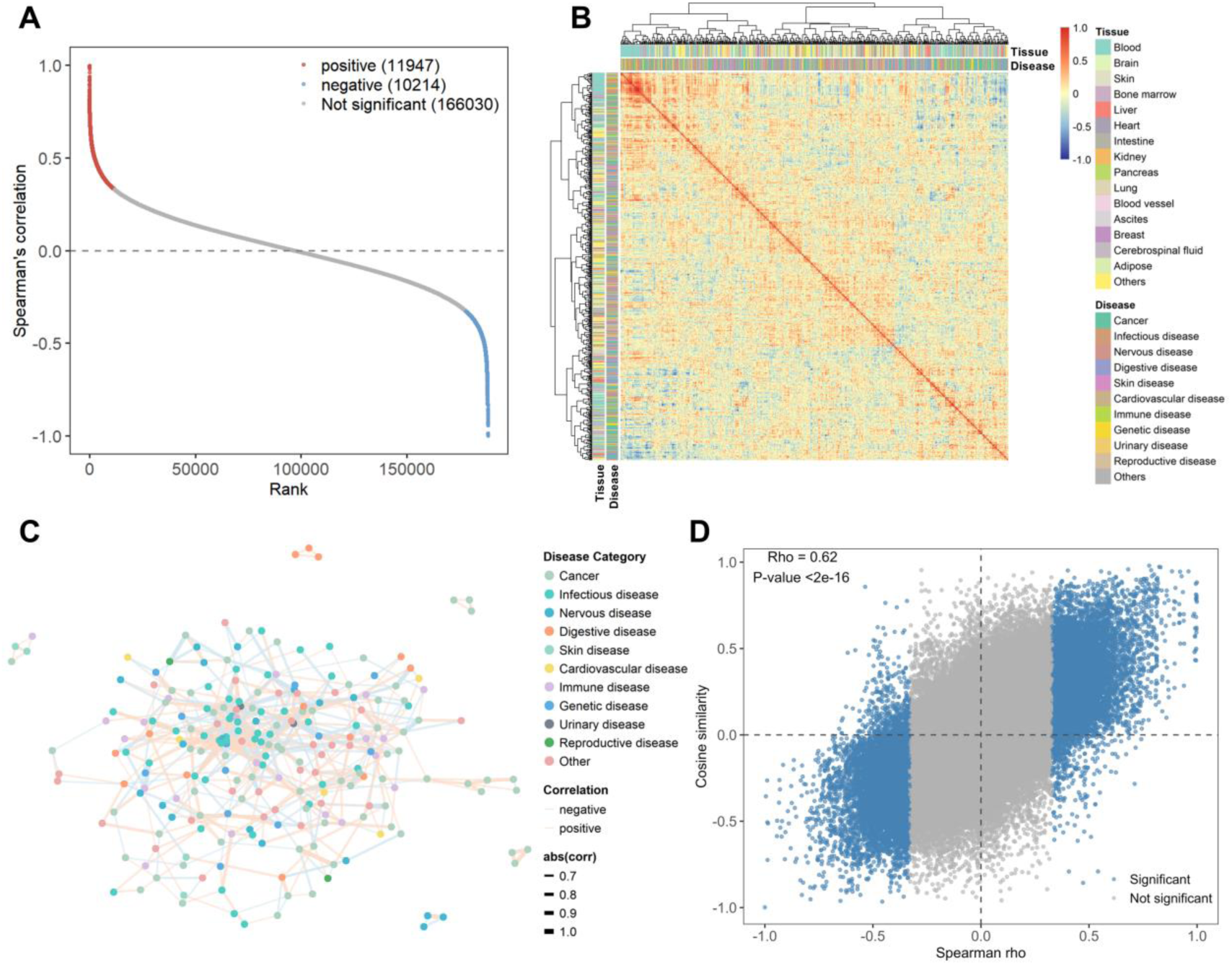
Identification of potential state bridges across biological states based on cell-composition change features. **(A)** Ranked distribution of Spearman correlation coefficients (Rho) among the 614 biological states. State bridges are ranked from the highest to the lowest correlation coefficient. Red indicates significant positive state bridges, blue indicates significant negative state bridges (P < 0.05). **(B)** Heatmap of Spearman correlations among biological states. Each row/column represents one state, and color indicates the Spearman correlation coefficient. Red and blue indicate positive and negative correlations, respectively. Annotations at the top and left indicate the tissue origin and disease category of each state. **(C)** State-association network constructed from Spearman correlations. State pairs with P < 0.05 and |Rho| > 0.65 were retained, and a k-core threshold of 2 was applied to extract the core network. Nodes represent biological states and are colored by disease category. Red and blue edges indicate positive and negative correlations, respectively, and edge width is proportional to |Rho|. **(D)** Concordance between Spearman correlation coefficients and cosine similarities. Each point represents a pair of biological states. Blue indicates state pairs with Spearman correlation P < 0.05.

Among these state bridges, we identified several relationships that are supported by previous studies and consistent with established biological knowledge. Five representative examples are shown in **Table 1**. For example, COVID-19 and tuberculosis showed a strong positive state bridge (Rho = 0.94, P = 2.44e-17). Alzheimer’s disease and Parkinson’s disease also exhibited a significant positive correlation (Rho = 0.92, P = 2.21e-15). Previous studies have reported biological links between these diseases at multiple levels, including immune responses, inflammatory processes, and neurodegenerative mechanisms^[21, 22]^. Recovery of these established relationships therefore supports the reliability of the state-bridge framework. In addition to positive bridges, we identified inverse relationships between disease states and established therapeutic interventions. For example, psoriasis was significantly negatively correlated with anti-IL-17A monoclonal antibody treatment (Rho = −0.66, P = 1.09e-5), suggesting that effective treatment may be accompanied by a cellular-remodeling pattern opposite to that observed in the disease state. Together, these representative relationships demonstrate that state bridges can capture both shared cellular-remodeling features across diseases and potential inverse remodeling relationships between diseases and therapeutic interventions.

**Table 1.** Five representative state bridges.

| **State A** | **State B** | **Spearman’s correlation** | **P value** |
|---|---|---|---|
| COVID-19 | Tuberculosis | 0.94 | 2.44e-17 |
| Alzheimer’s disease | Parkinson’s disease | 0.92 | 2.21e-15 |
| Dilated cardiomyopathy | Hypertrophic cardiomyopathy | 0.78 | 1.99e-8 |
| Psoriasis | Anti-IL-17A monoclonal antibody therapy | -0.66 | 1.09e-5 |
| Acute myeloid leukemia | Donor-derived natural killer cell infusion | -0.73 | 5.61e-7 |

We also used cosine similarity to assess the overall concordance in the direction of cellular changes between states. In total, 1,431 state pairs had a cosine similarity > 0.6, whereas 996 pairs had a cosine similarity < −0.6. Spearman correlation coefficients and cosine similarities were significantly positively correlated (Rho = 0.62, P < 2.00e-16), indicating overall consistency of cross-state relationships across different similarity measures (**Figure 5D**). We further applied Fisher’s exact test to the directions of cell-proportion changes, providing a complementary assessment of state relationships based on discrete directional patterns (Supplementary Table 2). Together, these complementary analyses consistently revealed widespread relationships in cellular-composition remodeling across states. The recovery of established biological relationships further supports the reliability of state bridges and provides a basis for identifying potential state relationships beyond existing knowledge.

### 3.5 Analysis of changed cell types in the context of biological states

We counted the number of changed cell types (NCC) among the 36 annotated cell types using a fold-change (FC) threshold of 1.5 (Supplementary Table 3). NCC varied considerably across biological states. For example, neuroblastoma (GSE137804) had the highest NCC, comprising 11 upregulated and 21 downregulated cell types, whereas Crohn’s disease (GSE298194) had only one downregulated cell type. Twelve biological states had an NCC greater than 25 (**Figure 6A**), and these were enriched in severe diseases such as neuroblastoma, glioblastoma, and pancreatic ductal adenocarcinoma, suggesting that such diseases may have a complex and comprehensive background of cell-type changes. Indeed, pancreatic ductal adenocarcinoma and glioblastoma are the two most malignant types of cancer. In contrast, 41 biological states had an NCC smaller than 5 (**Figure 6B**). Interestingly, many of these were immune diseases, such as graft-versus-host disease (GVHD), Crohn’s disease, Sjögren’s disease, inflammatory bowel disease, juvenile idiopathic arthritis, and myalgic encephalomyelitis.

**Figure 6.**
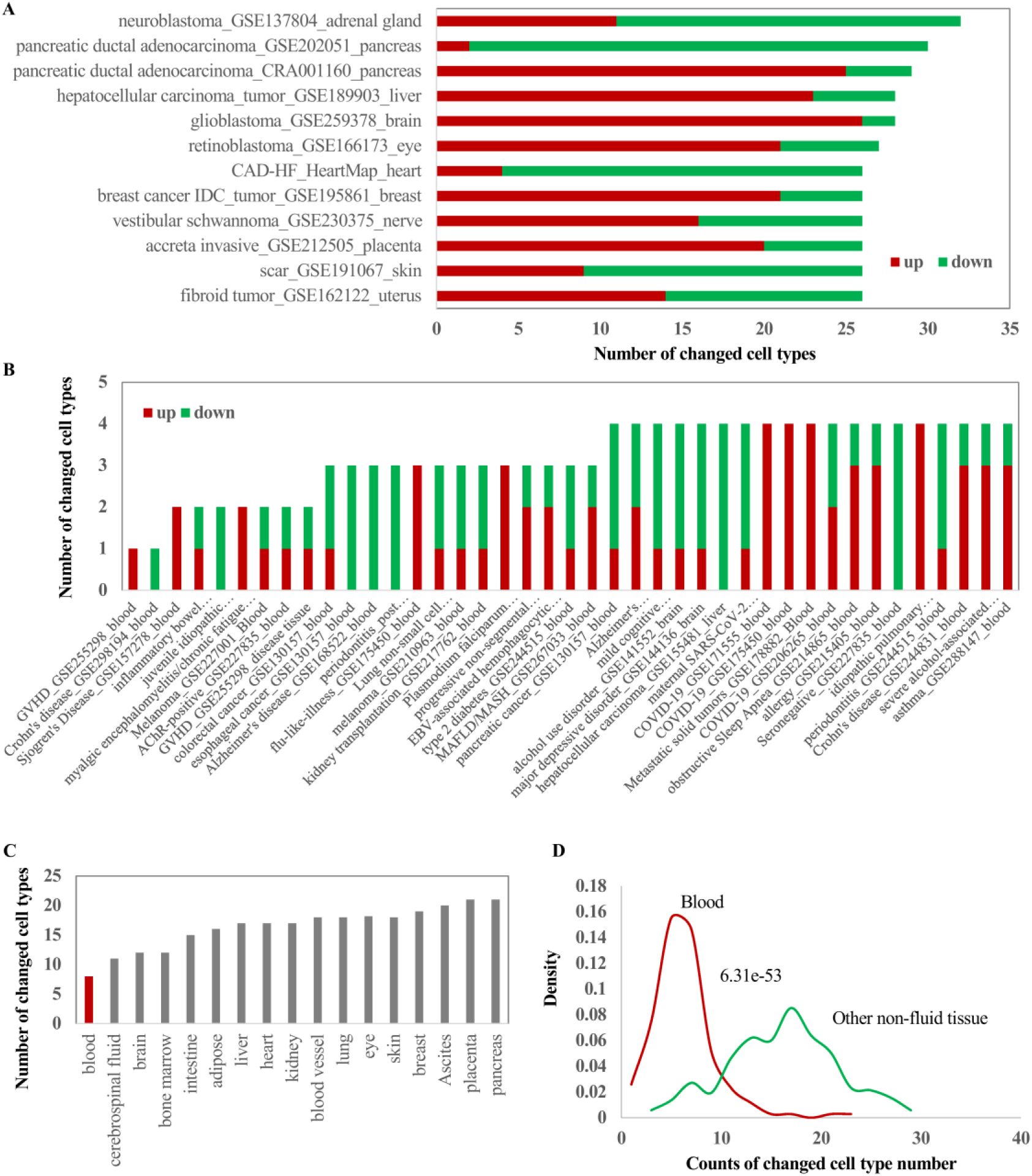
Analysis results of changed cell types in the context of biological states. **(A)** the 12 biological states with more than 25 changed cell types; **(B)** the 41 biological states with less than 5 changed cell types; **(C)** The number of changed cell types in corresponding tissue/organ origin for tissues/organs with at least five states; **(D)** The distributions of the changed cell type numbers of the blood-borne scRNA-seq data (red curve) and that of the other non-fluid tissue-based scRNA-seq data.

More interestingly, the above analysis revealed that biological states described by blood-borne scRNA-seq data tended to have a smaller NCC than those delineated by non-blood-derived scRNA-seq data. For example, none of the 12 states with the greatest NCC were derived from blood-borne scRNA-seq data, whereas 33 (80%) of the 41 states with the lowest NCC were (p = 4.72 × 10^−7^, Fisher’s exact test). We further grouped biological states by their corresponding tissue/organ of origin and averaged the NCC for tissues/organs with at least five states. As a result, states derived from blood had the smallest NCC (8), followed by cerebrospinal fluid (11), bone marrow (12), and brain (12) (**Figure 6C**). Interestingly, blood, cerebrospinal fluid, and bone marrow are all body fluids or fluid-like tissues. By comparison, placenta and pancreas were the top two tissues/organs with the greatest NCC (both 21) (**Figure 6C**), suggesting that state transitions in these tissues often involve complex and comprehensive changes in cell types. Comparison of blood-based and other non-fluid tissue-based states further indicated that the number of changed cell types in blood-based states was significantly smaller than in other non-fluid tissue-based states (median: 6 vs. 17, p = 6.31 × 10^−53^, **Figure 6D**). This finding suggests that, in the context of single-cell biology and medicine, blood may be far from reflecting the true pathological condition in diseased tissues/organs, and further implies that blood-based single-cell compositions may be inadequate for monitoring disease formation and progression, especially for early disease detection.

### 3.6 Shannon entropy reveals changes in cellular composition patterns across biological states

To further characterize the remodeling of cellular composition across biological states from the perspective of compositional evenness, we calculated the Shannon entropy based on the relative proportions of 36 annotated cell types in both case and control groups, and defined the entropy difference between the two groups as ΔH (Supplementary Table 1). Among the 614 biological states, 354 (57.7%) exhibited increased Shannon entropy, whereas 260 (42.3%) showed decreased entropy (**Figure 7A**). The magnitude of entropy changes varied considerably across biological states. For example, keloid (PRJNA844167) exhibited the largest increase in entropy (ΔH = 1.46), followed by gastric metaplasia (GSE134520, ΔH = 1.33) and autosomal dominant polycystic kidney disease (GSE185948, ΔH = 1.30). In contrast, a treatment-related comparison in pancreatic ductal adenocarcinoma (GSE202051) showed the greatest decrease in entropy ( Δ H = −1.93) (**Figure 7B**). Notably, a disease-associated state in another pancreatic ductal adenocarcinoma study (CRA001160) exhibited a marked increase in entropy (ΔH = 1.13), suggesting that disease states and therapeutic interventions may be associated with distinct directions of cellular compositional remodeling. Furthermore, the number of changed cell types (NCC) was positively correlated with the absolute entropy change (|ΔH|; Spearman’s ρ = 0.33, p = 4.68 × 10^−17^; **Figure 7C**), suggesting that more extensive cell-type changes are generally accompanied by greater alterations in cellular compositional evenness.

**Figure 7.**
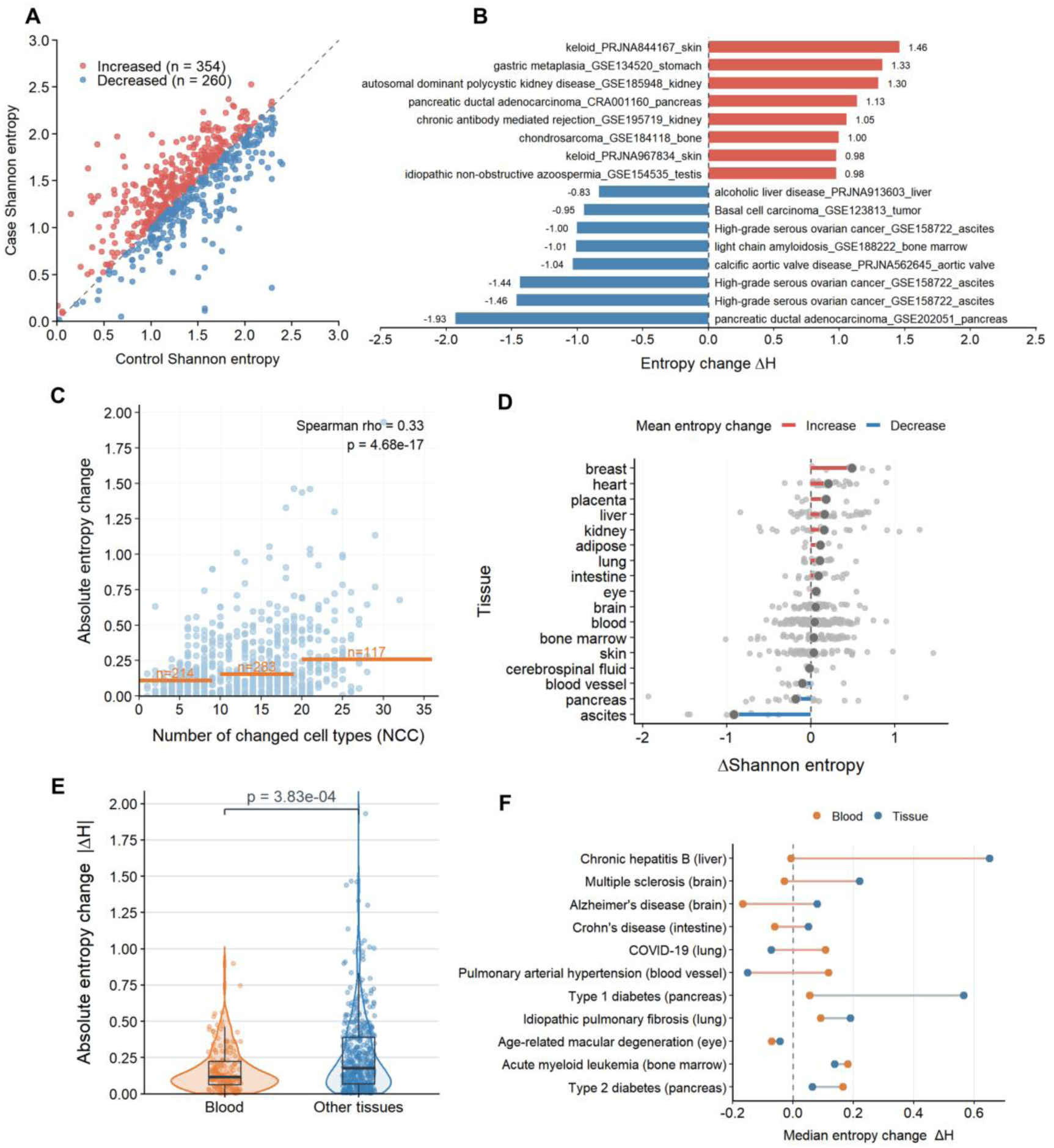
Shannon entropy analysis of cellular composition across biological states. **(A)** Comparison of Shannon entropy between case and control groups across 614 biological states. **(B)** The eight biological states with the largest increases and decreases in Shannon entropy (ΔH), respectively. **(C)** Correlation between the number of changed cell types (NCC) and the absolute entropy change (|ΔH|). Orange horizontal lines indicate the median |ΔH| within different NCC ranges. **(D)** Distribution of entropy changes across tissues/organs represented by at least five biological states. **(E)** Comparison of absolute entropy changes between blood-derived and other tissue-derived biological states. **(F)** Comparison of median entropy changes between blood and corresponding disease-associated tissues across 11 diseases.

We further grouped biological states according to their tissue of origin and calculated the mean ΔH for tissues represented by at least five states. The results revealed considerable variation in entropy changes across tissues (**Figure 7D**). Breast and heart derived states exhibited relatively large positive mean entropy changes (ΔH = 0.49 and 0.21, respectively), whereas pancreas- and ascites-derived states showed negative mean entropy changes (ΔH = −0.18 and −0.92, respectively). Notably, 30 of the 32 heart-derived states exhibited increased entropy. Interestingly, blood-derived states showed significantly smaller absolute entropy changes than states derived from other tissues (median |ΔH|: 0.12 vs. 0.18, p = 3.83 × 10^−4^, Mann-Whitney U test; **Figure 7E**), suggesting relatively limited changes in cellular compositional evenness in blood. Furthermore, comparisons between blood and corresponding diseased tissues across 11 diseases revealed that six exhibited opposite directions of entropy change (**Figure 7F**). For example, Alzheimer’s disease showed decreased entropy in blood but increased entropy in brain tissue, whereas COVID-19 exhibited the opposite pattern between blood and lung tissue. Together, these findings suggest that cellular compositional entropy remodeling is heterogeneous and tissue-dependent, and that changes in blood-derived cellular compositions may not fully reflect those occurring in diseased tissues.

## Discussion and Conclusion

This study establishes an analytical framework for systematically deciphering state bridges from scRNA-seq landscapes. Using the UCE model, we represented 614 biological states as 36-dimensional state vectors and mapped them into a unified cellular-composition remodeling space. We ultimately identified 22,161 statistically significant bridges among the 614 states (P < 0.05). These included many relationships already supported by existing knowledge. More importantly, we also uncovered previously unrecognized state bridges, including the bridge between xx and xx, which was further validated in animal experiments. These findings suggest that the value of large-scale scRNA-seq data extends beyond separately describing changes in cellular composition within individual states. By transforming these otherwise isolated changes into comparable state-level features, our framework enables potential biological relationships beyond existing knowledge to be identified. In this way, our study provides a new perspective for investigating disease associations and potential intervention relationships.

Our findings further suggest that relationships among biological states are not restricted to molecular expression patterns but can also be reflected in cellular composition. Gene-expression-based state representations, such as the Connectivity Map (CMap)^[6]^, CREEDS^[8]^, LINCS^[9]^ and iLINCS^[10]^, can effectively capture changes in intracellular transcriptional programs. However, these molecular signatures primarily describe responses within cells and do not directly represent changes in the relative abundance of different cell populations. This additional layer of information is important. Disease progression and therapeutic intervention are often accompanied by coordinated changes across multiple cell populations, including immune-cell recruitment, stromal-cell expansion and parenchymal-cell depletion^[2, 23]^. Together, these changes shape the tissue microenvironment and its functional state. Changes in cellular composition therefore provide a direct representation of this reorganization of cellular ecology and capture patterns of cellular remodeling across biological states. In this sense, a cell-composition signature does more than add another feature type to existing signature-matching approaches. It provides a distinct level of state representation that directly characterizes the reorganization of cellular ecology.

More importantly, state relationships defined by cellular-composition remodeling may provide biological insight beyond statistical association alone. Positive bridges indicate convergent changes across multiple cell populations and may therefore point to shared pathological processes, changes in the tissue microenvironment or multicellular response mechanisms. The recovery of established disease relationships in our analysis supports this interpretation. For example, Alzheimer’s disease and Parkinson’s disease showed a significant positive state bridge (Rho = 0.92, P = 2.21e-15). The relationship between these two neurodegenerative diseases has been extensively studied, with previous work focusing mainly on molecular and pathological features such as pathological proteins, neuroinflammation, mitochondrial dysfunction and neurodegenerative processes^[22, 24, 25]^. In contrast, our framework recovered this relationship solely from patterns of cellular-composition change. This finding suggests that shared features between diseases may not be confined to intracellular molecular programs but can also manifest at a higher level as convergent remodeling of cellular ecology.

In contrast, inverse state bridges provide another dimension of information with potential translational relevance. If a disease state represents a cellular-remodeling pattern that deviates from homeostasis, an effective intervention may shift this remodeling in the opposite direction. Thus, when a disease state and a treatment state show opposing changes across multiple cell populations, this negative association may indicate a potential reversal of cellular-composition remodeling. In our analysis, psoriasis showed a significant negative correlation with anti-IL-17A monoclonal antibody treatment (Rho = −0.66, P = 1.09e-5). IL-17A is a key effector pathway in psoriatic inflammation, and monoclonal antibodies targeting IL-17A are widely used for the treatment of psoriasis^[26, 27]^. Notably, our observation extends beyond the established correspondence between a therapeutic target and disease mechanism. Instead, the disease and treatment states exhibited opposing patterns of cellular-composition change at the global level. This suggests that therapeutic effects may not be limited to suppression of specific inflammatory pathways but may also be reflected in tissue-level reorganization of cellular ecology, partially shifting disease-associated multicellular composition changes in the opposite direction.

It is important to emphasize that a state bridge is primarily a relationship-discovery tool rather than a causal-inference framework. Similarity between two states in the 36-dimensional cellular-composition space does not imply that they share the same etiology or molecular mechanisms. Likewise, inverse remodeling between a disease state and a treatment state does not by itself establish that the intervention would necessarily be effective against another disease. State bridges are therefore better viewed as a hypothesis-generating framework. They can first be used to prioritize potentially informative relationships within a large-scale state space, after which their biological basis can be examined through transcriptional programs, pathway activity, cell–cell communication, and experimental validation. In the future, integrating cellular-composition changes with intracellular molecular states and multi-omics information could further improve the resolution and mechanistic interpretability of state bridges.

## Data availability

The raw data used in our study are publicly available. All data supporting the findings of this study are included in the article and supplementary materials.

## Acknowledgements

We than lab members for valuable discussions.

## Authors’ contributions

Q.C. proposed the idea. Q.C. and B.K. supervised the study. Q.T. performed the study. M.L. and C.C. took part in the discussion. Q.T. wrote the draft manuscript and Q.C. edited the manuscript.

## References

[1] Sikkema, L., et al., An integrated cell atlas of the lung in health and disease. Nat Med, 2023. 29(6): p. 1563–1577.

[2] Thomas, T., et al., A longitudinal single-cell atlas of anti-tumour necrosis factor treatment in inflammatory bowel disease. Nat Immunol, 2024. 25(11): p. 2152–2165.

[3] Friedrich, M., et al., IL-1-driven stromal-neutrophil interactions define a subset of patients with inflammatory bowel disease that does not respond to therapies. Nat Med, 2021. 27(11): p. 1970–1981.

[4] Bhattacharya, M. and P. Ramachandran, Immunology of human fibrosis. Nat Immunol, 2023. 24(9): p. 1423–1433.

[5] Chu, Y., et al., Pan-cancer T cell atlas links a cellular stress response state to immunotherapy resistance. Nat Med, 2023. 29(6): p. 1550–1562.

[6] Lamb, J., et al., The Connectivity Map: using gene-expression signatures to connect small molecules, genes, and disease. Science, 2006. 313(5795): p. 1929–35.

[7] Sirota, M., et al., Discovery and preclinical validation of drug indications using compendia of public gene expression data. Sci Transl Med, 2011. 3(96): p. 96-77.

[8] Wang, Z., et al., Extraction and analysis of signatures from the Gene Expression Omnibus by the crowd. Nat Commun, 2016. 7: p. 12846.

[9] Subramanian, A., et al., A Next Generation Connectivity Map: L1000 Platform and the First 1,000,000 Profiles. Cell, 2017. 171(6): p. 1437–1452.e17.

[10] Pilarczyk, M., et al., Connecting omics signatures and revealing biological mechanisms with iLINCS. Nat Commun, 2022. 13(1): p. 4678.

[11] Luecken, M.D., et al., Benchmarking atlas-level data integration in single-cell genomics. Nat Methods, 2022. 19(1): p. 41–50.

[12] Xu, C., et al., Automatic cell-type harmonization and integration across Human Cell Atlas datasets. Cell, 2023. 186(26): p. 5876–5891.e20.

[13] Rosen, Y., et al., Universal cell embedding provides a foundation model for cell biology. Nature, 2026. 656(8126): p. 183–191.

[14] Li, M., et al., Rediscovering publicly available single-cell data with the DISCO platform. Nucleic Acids Res, 2025. 53(D1): p. D932–d938.

[15] Ma, S., et al., scCT-DB, an omnibus for cancer patient-derived paired pre- and post-treatment single-cell transcriptomes to reveal drug perturbation and drug resistance mechanisms. Nucleic Acids Res, 2026. 54(D1): p. D1616–d1633.

[16] Datar, Y., et al., An integrated cell atlas of 2.4 million cardiac cells across 209 individuals in health and disease. Nat Cardiovasc Res, 2026. 5(7): p. 624–637.

[17] Wang, J.C., et al., Cerebrovascular vulnerability and fibrosis in human brain aneurysms. Nat Neurosci, 2026. 29(8): p. 1814–1825.

[18] Wolf, F.A., P. Angerer, and F.J. Theis, SCANPY: large-scale single-cell gene expression data analysis. Genome Biol, 2018. 19(1): p. 15.

[19] Wolock, S.L., R. Lopez, and A.M. Klein, Scrublet: Computational Identification of Cell Doublets in Single-Cell Transcriptomic Data. Cell Syst, 2019. 8(4): p. 281–291.e9.

[20] Jones, R.C., et al., The Tabula Sapiens: A multiple-organ, single-cell transcriptomic atlas of humans. Science, 2022. 376(6594): p. eabl4896.

[21] Najafi-Fard, S., et al., Characterization of the immune impairment of patients with tuberculosis and COVID-19 coinfection. Int J Infect Dis, 2023. 130 **Suppl 1**: p. S34–s42.

[22] Weiner, H.L., Immune mechanisms and shared immune targets in neurodegenerative diseases. Nat Rev Neurol, 2025. 21(2): p. 67–85.

[23] de Visser, K.E. and J.A. Joyce, The evolving tumor microenvironment: From cancer initiation to metastatic outgrowth. Cancer Cell, 2023. 41(3): p. 374–403.

[24] Wilson, D.M., 3rd, et al., *Hallmarks of neurodegenerative diseases*. Cell, 2023. 186(4): p. 693–714.

[25] Patani, R., G.E. Hardingham, and S.A. Liddelow, Functional roles of reactive astrocytes in neuroinflammation and neurodegeneration. Nat Rev Neurol, 2023. 19(7): p. 395–409.

[26] Ghoreschi, K., et al., Therapeutics targeting the IL-23 and IL-17 pathway in psoriasis. Lancet, 2021. 397(10275): p. 754–766.

[27] Mills, K.H.G., IL-17 and IL-17-producing cells in protection versus pathology. Nat Rev Immunol, 2023. 23(1): p. 38–54.

